# Centromere-specifying nucleosomes persist in aging mouse oocytes in the absence of nascent assembly

**DOI:** 10.1101/2023.05.18.541332

**Authors:** Arunika Das, Katelyn G. Boese, Kikue Tachibana, Sung Hee Baek, Michael A. Lampson, Ben E. Black

## Abstract

**SUMMARY:** Centromeres direct genetic inheritance but are not themselves genetically encoded. Instead, centromeres are defined epigenetically by the presence of a histone H3 variant, CENP-A^1^. In cultured somatic cells, an established paradigm of cell cycle-coupled propagation maintains centromere identity: CENP-A is partitioned between sisters during replication and replenished by new assembly, which is restricted to G1. The mammalian female germline challenges this model because of the cell cycle arrest between pre-meiotic S-phase and the subsequent G1, which can last for the entire reproductive lifespan (months to decades). New CENP-A chromatin assembly maintains centromeres during prophase I in worm and starfish oocyte^2,3^, suggesting that a similar process may be required for centromere inheritance in mammals. However, we show that centromere chromatin is maintained long-term independent of new assembly during the extended prophase I arrest in mouse oocytes. Conditional knockout of Mis18α, an essential component of the assembly machinery, in the female germline at the time of birth has almost no impact on centromeric CENP-A nucleosome abundance nor any detectable detriment to fertility.

## RESULTS AND DISCUSSION

In contrast to genetic information encoded in our genome, minimal epigenetic information is inherited because most parental epigenetic marks are removed and reprogrammed in germ cells and the early embryo^4,5^. A key exception is the centromeric histone H3 variant, CENP-A^6–8^. Although centromeres direct the process of genetic inheritance by connecting chromosomes to spindle microtubules, they are not encoded in DNA, but rather epigenetically specified by nucleosomes containing CENP-A^9^. Thus, CENP-A nucleosomes are inherited to preserve centromere identity.

Studies in cycling somatic cells have established a general paradigm for propagation of epigenetic information between cell cycles. DNA or histone modifications are partitioned between sister chromatids during DNA replication, and then replenished by “reader” proteins that recognize the modification and “writers” that extend it to adjacent nucleosomes^10^. CENP-A, follows this paradigm, with new assembly restricted to G1 by CDK1/2 activity^11–13^. CENP-A and its bound partner histone H4 molecules are remarkably stable in tissue culture cells^14,15^, with measured CENP-A turnover rates explained entirely through the dilution when existing CENP-A is partitioned to replicated centromeric DNA during S-phase each cell cycle^14^. This paradigm poses a challenge in the mammalian female germline because of the extended prophase I cell cycle arrest, after replication but before an opportunity for new G1 assembly, which can last for the entire reproductive lifespan of the animal^16^. Centromeres are preserved throughout the arrest (>1 year in mouse) in the absence of new *Cenpa* transcription, as shown by conditional knockout^17^ of the *Cenpa* gene (Figure 1A). CENP-A nucleosomes are therefore either replenished by new assembly in contrast to the somatic cell paradigm, drawing on a stable pool of CENP-A protein, or stable for the entire duration of the arrest (Figure 1B).

**Figure 1:**
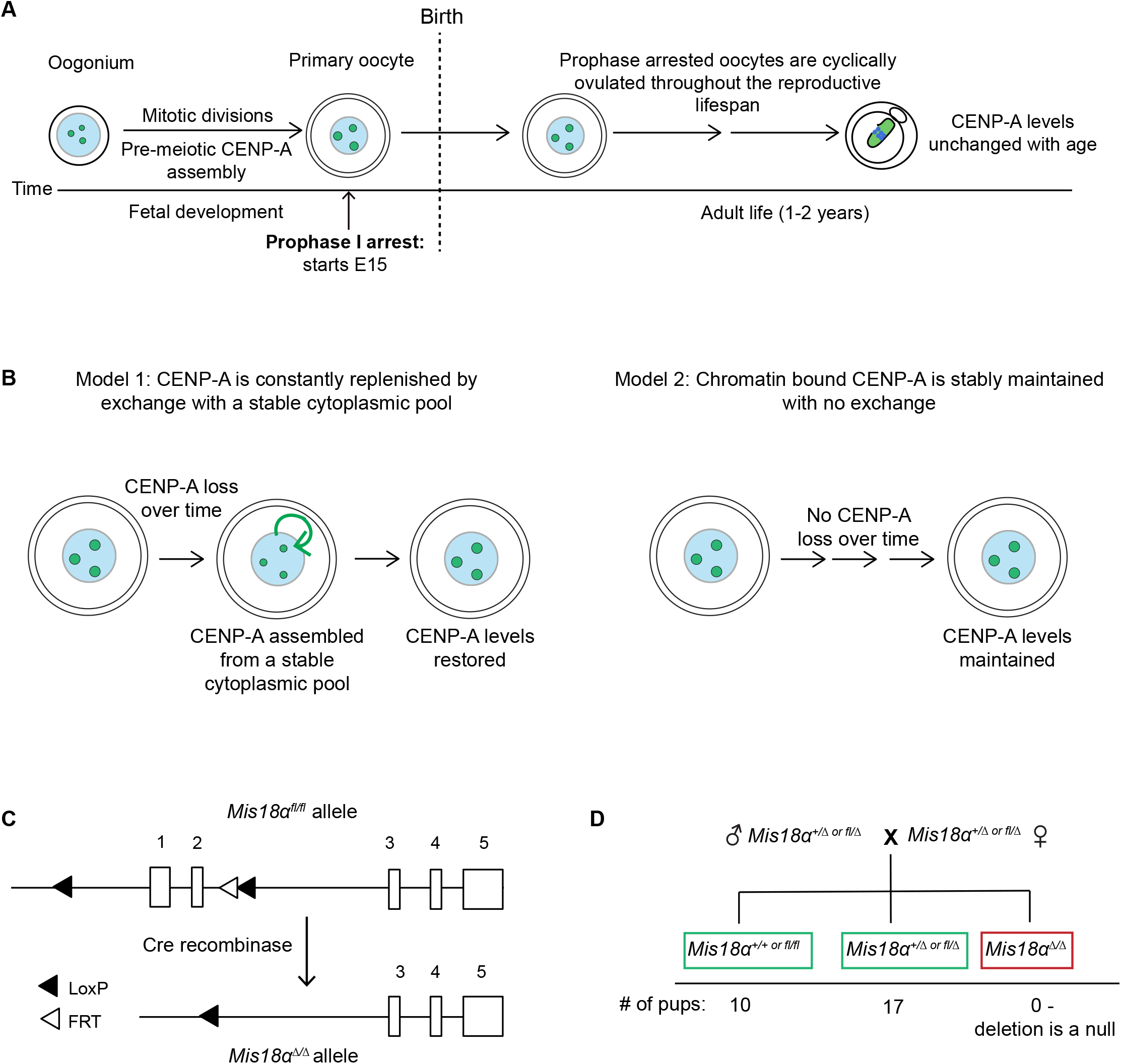
Testing models for stable retention of CENP-A chromatin in oocytes. (A) Schematic showing maintenance of CENP-A chromatin across the oogenesis timeline. Oocytes arrested in prophase I are cyclically recruited to begin growth before ovulation and maturation. Prophase arrest prior to recruitment can last up to 2 years in mice. Centromeric CENP-A levels are stable over time even if the *Cenpa* gene is eliminated shortly after birth^17^. (B) Models for retention of CENP-A in meiotic prophase I arrested eggs. (C) Schematic of the *Mis18α* conditional knockout gene locus^25^. The 1^st^ and 2^nd^ *Mis18α* protein coding exons are flanked by LoxP sites (floxed, fl). The FRT site is a remnant from FLP mediated excision of the Neomycin cassette in the original construct used to generate the KO animals. (D) Genotype frequencies of the progeny from a cross of *Mis18α*^*+/Δ or fl/Δ*^ heterozygotes (See Figure S2 for generation of heterozygotes). Number of litters = 6, number of pups = 27.

Studies in worm and starfish oocytes show nascent CENP-A chromatin assembly in prophase I (akin to a G2 biochemical cell cycle state in a somatic cell)^2,3^. In mouse, continual deposition of nucleosomes containing another H3 variant, H3.3, during oocyte development is required for oocyte genome integrity and, ultimately, for fertility^18^. These studies suggest that new assembly may maintain CENP-A chromatin through the prophase I arrest. To test this prediction, we created an oocyte specific conditional knockout (cKO) of an essential component of the CENP-A deposition machinery, Mis18α (Mis18a in mouse but referred to as Mis18α for simplicity)^19,20^, to prevent nascent CENP-A chromatin assembly. Mis18α is part of the Mis18 complex, which recruits the CENP-A chaperone, HJURP, bound to nascent CENP-A/histone H4 dimers^21–23^, to centromeres. The Mis18 complex is required for both nascent CENP-A chromatin assembly in G1 and replication-coupled CENP-A chromatin assembly in S phase in cycling somatic cells^24^. Specifically relevant to our test of ongoing chromatin assembly in the mouse oocyte, prophase I assembly in both worms and starfish oocytes requires the Mis18 complex^2,3^. Thus, If CENP-A chromatin is replenished by new assembly, CENP-A nucleosome levels would decay in Mis18α knockout oocytes.

For conditional knockout, we used a floxed *Mis18α* allele in which the first two exons, encoding the YIPPEE domain necessary for CENP-A deposition, are flanked by *LoxP* sites^25^ (Figure 1C, S1). To confirm that Cre-mediated excision generates a null allele, we crossed *Mis18α* heterozygous parents and did not recover any progeny homozygous for the deletion, as expected because *Mis18α* is an essential gene^25^ (Figures 1D, S2). Combining the floxed allele with *Cre* recombinase driven by oocyte specific promoters, we generated cKOs of *Mis18α* either early or late in prophase I (Figure S2). The early *Cre* driver (*Gdf9-Cre*) deletes *Mis18α* in arrested oocytes 2 days post birth, preventing assembly of new CENP-A nucleosomes for nearly the entire lifespan of the animal^26^. The late *Cre* driver (*Zp3-Cre*) deletes the gene during oocyte growth, 2-3 weeks prior to ovulation^26^ (Figure 2A).

**Figure 2:**
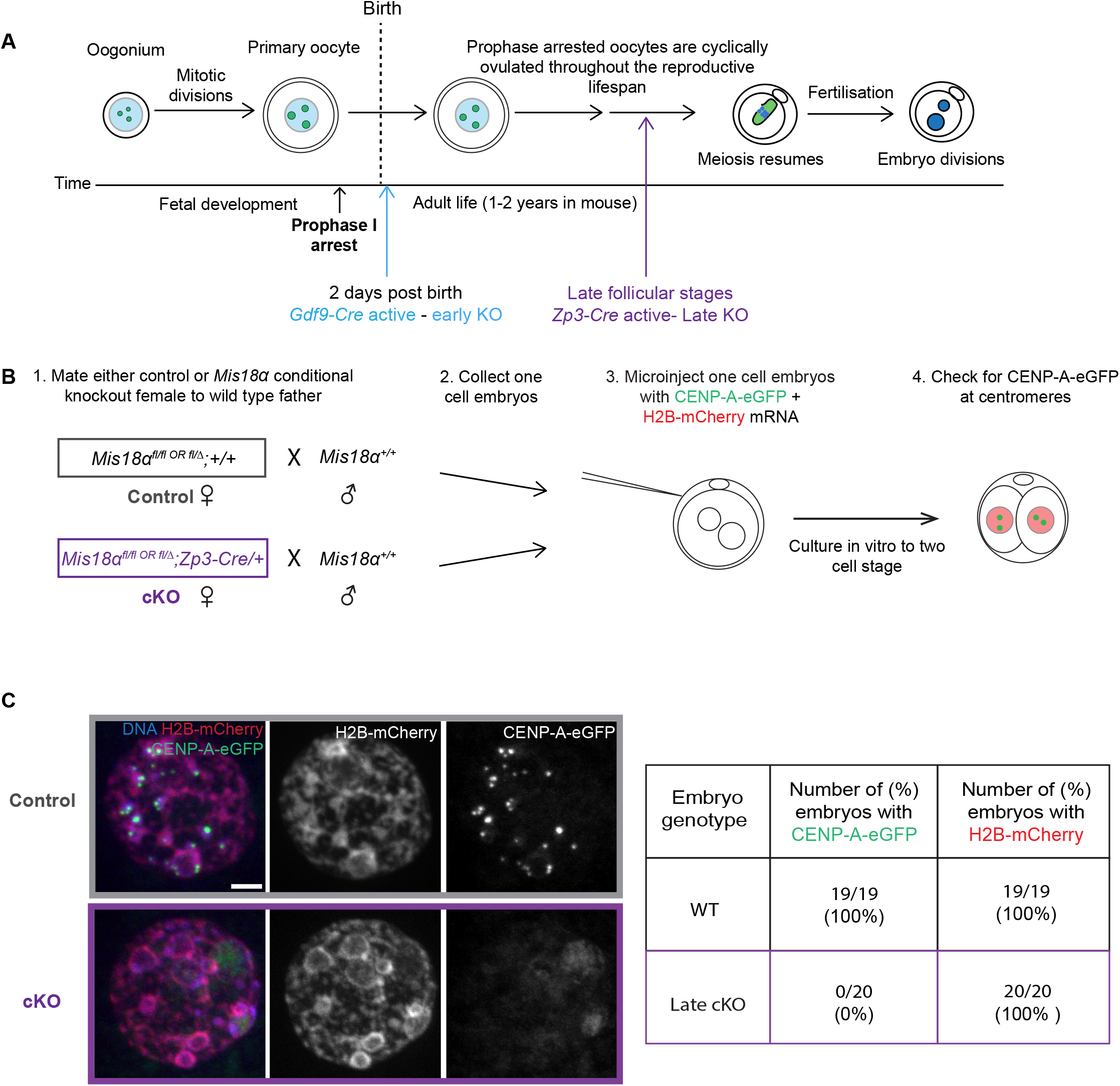
Maternally deposited Mis18α protein is eliminated in late KO oocytes. (A) Schematic showing early and late *Mis18α* knockout depending on the timing of *Cre* recombinase expression driven by either *Gdf9* or *Zp3* promoters (colored arrows). (B) Experimental design to test for a stable pool of Mis18α protein in the maternal cytoplasm. CENP-A-eGFP and H2B-mCherry mRNA are injected into one cell embryos, and CENP-A foci are assessed after the first embryonic mitosis when assembly is expected to occur but before zygotic genome activation. (C) Images show H2B-mCherry, CENP-A-eGFP, and DNA (DAPI) in interphase two cell embryos from control or cKO mothers. Table shows frequencies of detectable assembly for CENP-A-eGFP and H2B-mCherry, obtained from 2 independent matings. Scale bars = 5 μm.

As a functional assay for *Mis18α* deletion, we tested for CENP-A chromatin assembly in embryos from late KO mothers (driven by *Zp3-Cre*) and a wild type father (Figure 2B). Prior to zygotic genome activation at the two-cell stage, CENP-A deposition during the early embryonic mitotic cycles depends solely on maternal Mis18α protein. To assess new chromatin assembly, we injected mRNAs encoding CENP-A-eGFP and H2B-mCherry into one-cell embryos derived from either KO mothers or wild type mothers as controls (Figure 2B). H2B-mCherry serves as a positive control for chromatin assembly because it utilizes a deposition pathway distinct from that of CENP-A and therefore does not require the Mis18 complex. Based on the paradigm established in cycling somatic cells, we expected new CENP-A chromatin assembly in G1 after the first embryonic mitosis. Indeed, CENP-A-eGFP localized to centromeres in all control embryos (100%, N = 19) from wild type mothers. In contrast, none of the embryos from late *Mis18α* KO mothers (0%, N = 20) contained detectable centromeric CENP-A-eGFP (Figure 2C). H2B-mCherry was present in chromatin in 100% of both control and KO injected embryos as expected. This finding establishes that even after a relatively short duration KO, *Mis18α* deletion in oocytes abrogates nascent CENP-A chromatin assembly in early embryos.

Next, we measured fertility of *Mis18α* KO mothers as the ultimate test of centromere function, because centromeres are required for the meiotic divisions and for maternal centromere inheritance, and even partial reduction of maternal CENP-A chromatin lowers fertility^8^. All *Mis18α* early and late KO mothers were fertile when crossed to wild type males (Figure 3A). We also confirmed that both *Cre* recombinases have a 100% deletion efficiency as we did not recover a floxed allele of *Mis18α* inherited from a cKO mother (Figure 3B). Combined with our embryo experiments, this result confirms that every oocyte with a combination of floxed alleles and *Cre* lacks *Mis18α* protein and that fertility is not a consequence of inefficient *Cre* activity in some oocytes.

**Figure 3:**
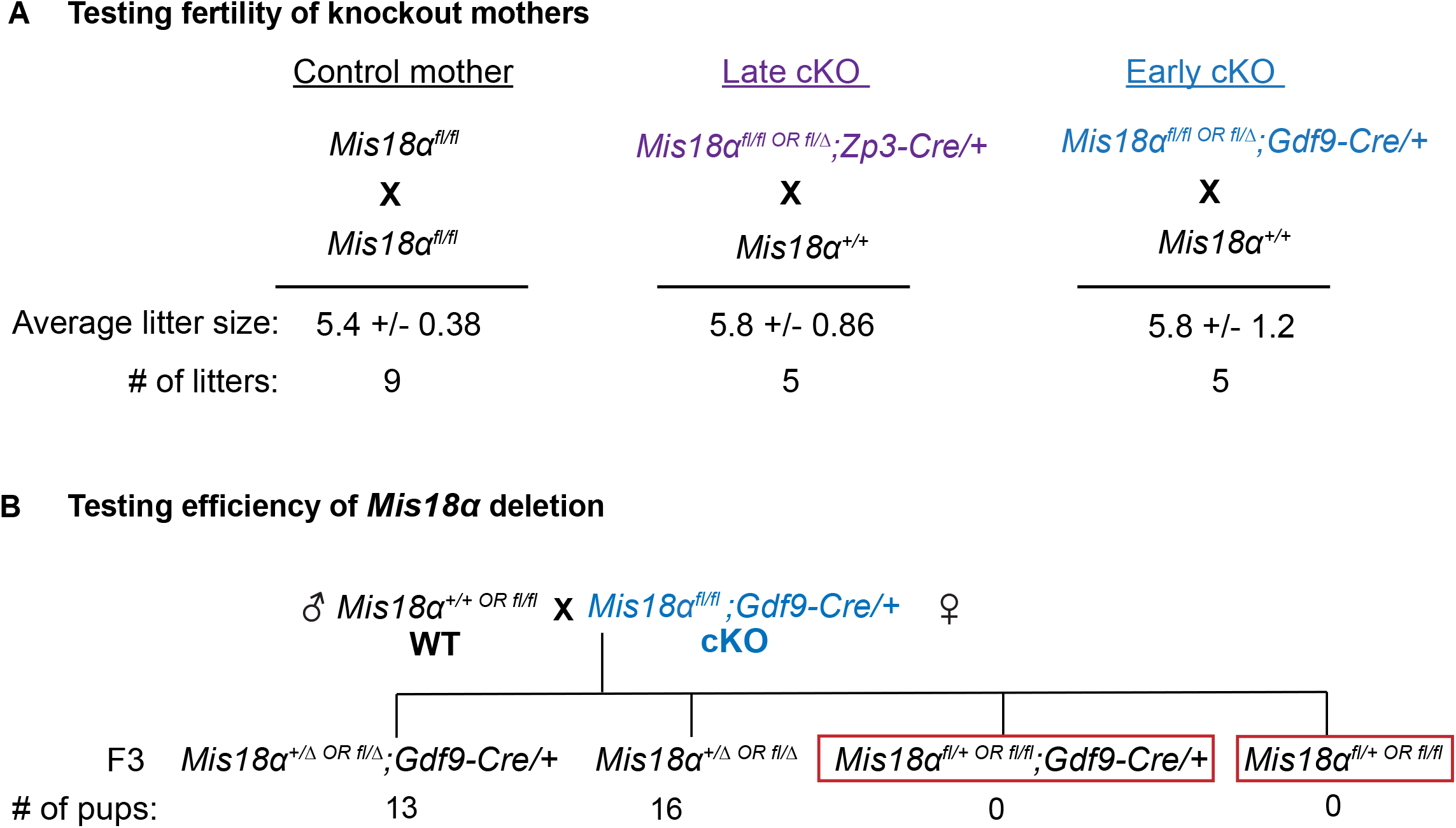
*Mis18α* knockout mothers can support fertility. (A) Litter sizes for control, early cKO, and late cKO mothers (age 2-4 months) crossed to wild type males. (B) Genotype frequencies of the progeny from a cross between a WT father and a cKO mother (N= 29 pups, 5 litters).

Even though the cKO mice are fertile, there could still be a reduction in CENP-A nucleosomes over time. Therefore, we tested if CENP-A levels decay in the late KO oocytes. We predict that if CENP-A nucleosomes are continually replenished by new assembly from a stable pool of CENP-A protein, centromeric CENP-A levels would decline in the KO oocytes compared to control oocytes. We did not see any reduction in centromeric CENP-A levels in late KO oocytes compared to controls (Figures 4A, B).However, *Mis18α* is deleted for only 2-3 weeks before ovulation in the late KO, leaving open the possibility of a more dramatic reduction in CENP-A nucleosomes on longer timescales.

**Figure 4:**
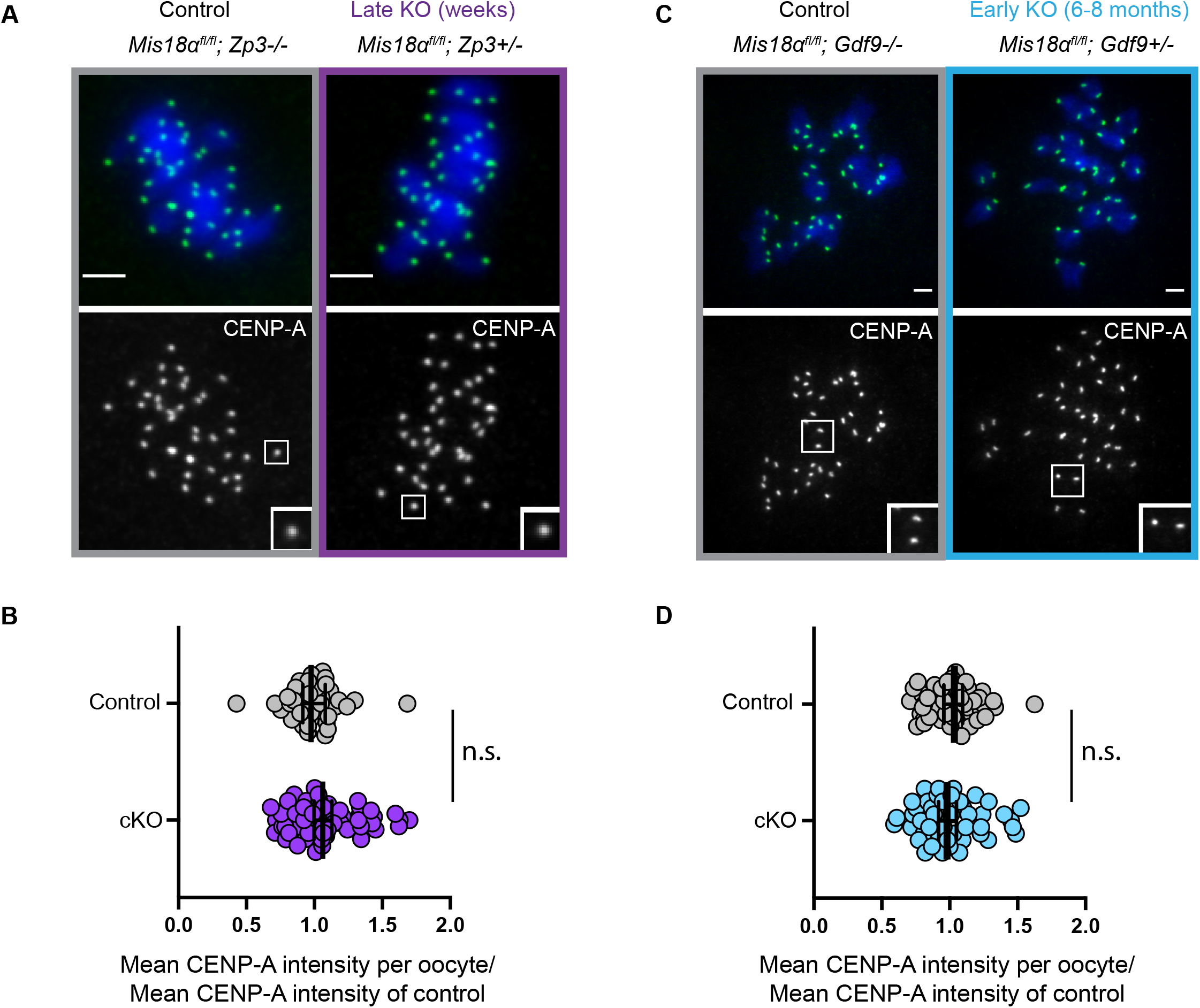
Pre-meiotically assembled CENP-A chromatin is maintained in prophase arrested oocytes without new assembly. (A, C) Images show metaphase I oocytes stained for CENP-A (green) and DNA (DAPI, blue) from late cKO (A) or early cKO (B) mothers. Scale bars = 5 μm. (B, D) Quantifications of CENP-A intensities in metaphase I oocytes from late cKO (B) or early cKO (D) mothers. Each data point represents average CENP-A levels at centromeres in each oocyte normalized to the control mean. Late cKO (*Zp3-Cre*, purple): mean ± S.E.M. = 1.1 ± 0.034 compared to control, number of oocytes = 58, number of centromeres = 1512, N = 7 females) and *Cre* negative control oocytes (number of oocytes = 40, number of centromeres = 962, N = 5 females). Early cKO (*Gdf9-Cre*, blue): mean ± S.E.M. = 1.0 ± 0.068 (number of oocytes = 47, number of centromeres = 1746, N = 7 females) relative to control (number of oocytes = 50, number of centromeres = 2019, N = 8 females); n.s.: Mann-Whitney U Test. Error bars: geometric mean with 95% confidence interval. The early cKO mothers are aged 6-8 months, and their oocytes lack Mis18α for the entire lifespan.

Thus, we leveraged the early KO (*Gdf9-Cre*) oocytes that delete *Mis18α* shortly after birth, to test if centromeric CENP-A levels decline after months without new assembly (Figure 4C, D). We aged control and early KO females for 6-8 months and found that CENP-A levels were not significantly reduced compared to the controls. Although not statistically significant, we measured a ∼3% decrease in CENP-A levels in the KO oocytes relative to control (Figure 4D) that equates to loss at a rate of ∼0.02% per day over 180 days. With ∼200 CENP-A nucleosomes estimated at each mammalian centromere^27^, any such loss in signal would be in the range of one nucleosome per month. In comparison, new assembly in starfish oocytes is estimated at a rate of 2% per day, based on centromere localization of GFP-tagged exogenous CENP-A in cells cultured *in vitro* for 10 days^3^.

## CONCLUSION

In conclusion, these results provide clear evidence supporting long term retention of pre-meiotically assembled CENP-A nucleosomes as the dominant pathway for maintaining centromere identity in mammalian oocytes. This is in stark contrast to H3.3, whose ongoing deposition into bulk chromatin during prophase I is essential for normal oocyte chromatin structure and fertility^18^. In addition, our findings have implications for human female meiosis, which is inherently error prone and especially vulnerable to aging. With advancing maternal age at childbirth, mechanisms that preserve centromeres in aging oocytes gain increasing significance. Our findings can now direct future research into the mechanisms that underlie CENP-A retention in mammalian oocytes. Previous studies of centromere chromatin suggest multiple potential molecular mechanisms that could contribute to its stability: the relatively low level of transcription of centromeric DNA^28–31^ relative to genic regions harboring histone H3.3 nucleosomes^32^; structural features that differ from canonical nucleosomes, including internal structural rigidity at the CENP-A/histone H4 interface^33,34^; and non-histone CCAN components that bind and stabilize CENP-A nucleosomes in tissue culture cells^14,15,35^. In sum, it remains to be seen if the mechanisms that function to retain CENP-A chromatin over short periods of time in cycling cells also contribute to extreme stability during oogenesis.

## ACKNOWLEDGEMENTS

We would like to thank M.T. Levine and R. M. Schultz for feedback on the manuscript. This work was supported by the National Institute of Child Health and Development grant HD058730 (M.A.L. and B.E.B.)

## AUTHOR CONTRIBUTIONS

A.D., M.A.L., and B.E.B. designed the project and wrote this manuscript with input from co-authors. M.A.L. and B.E.B. provided supervision and sourced funding. M.A.L. and B.E.B. and K.T. conceived the project. A.D. performed all the experiments and analyzed the data. A.D. and K.G.B. maintained mouse colony, genotyping, and quantified the data. S.B. and K.T. provided the *Mis18α* knockout mouse line and K.T. did initial experimentation.

## INCLUSION AND DIVERSITY

We support inclusive, diverse, and equitable conduct of research.

## STAR METHODS

### Key resources table

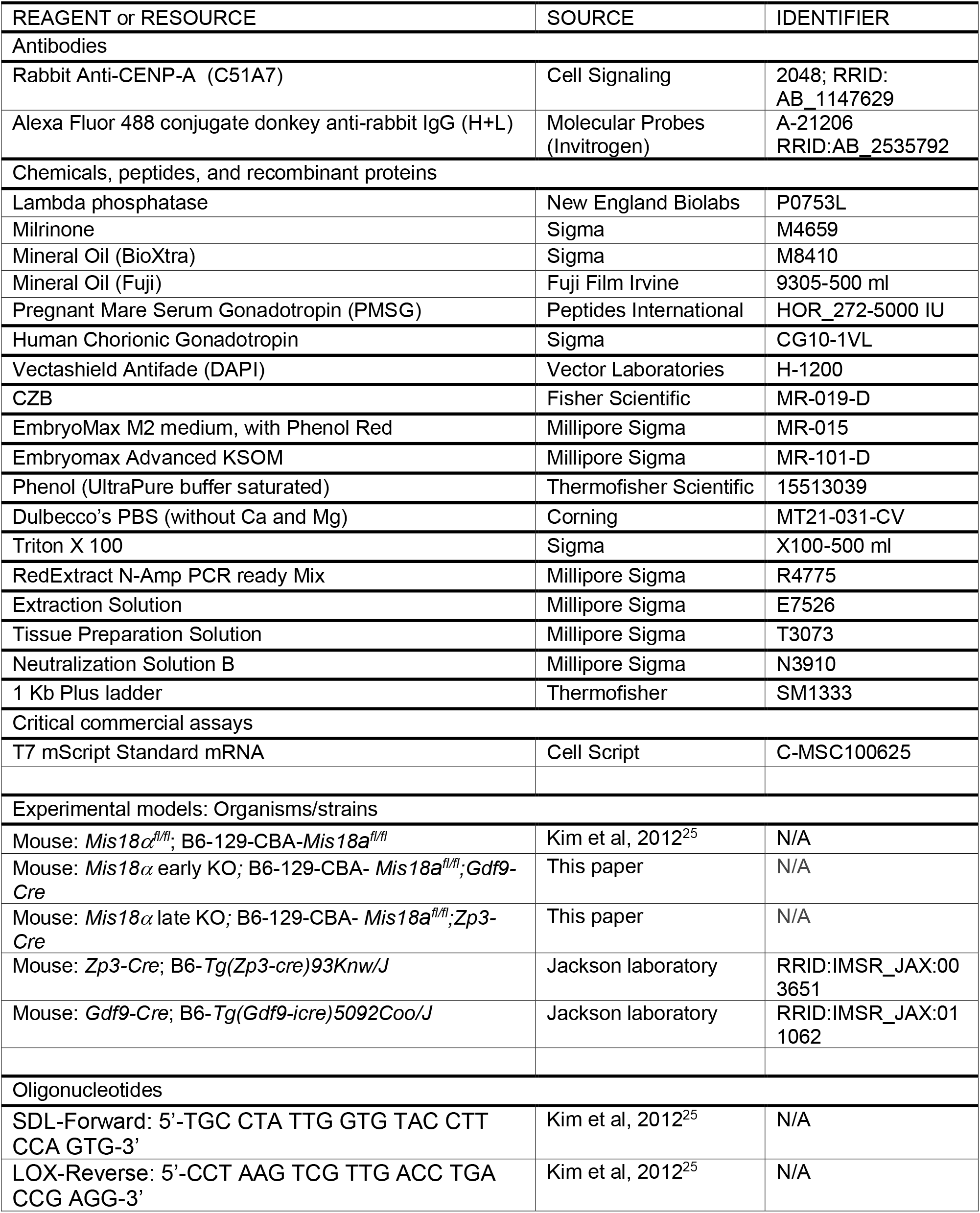
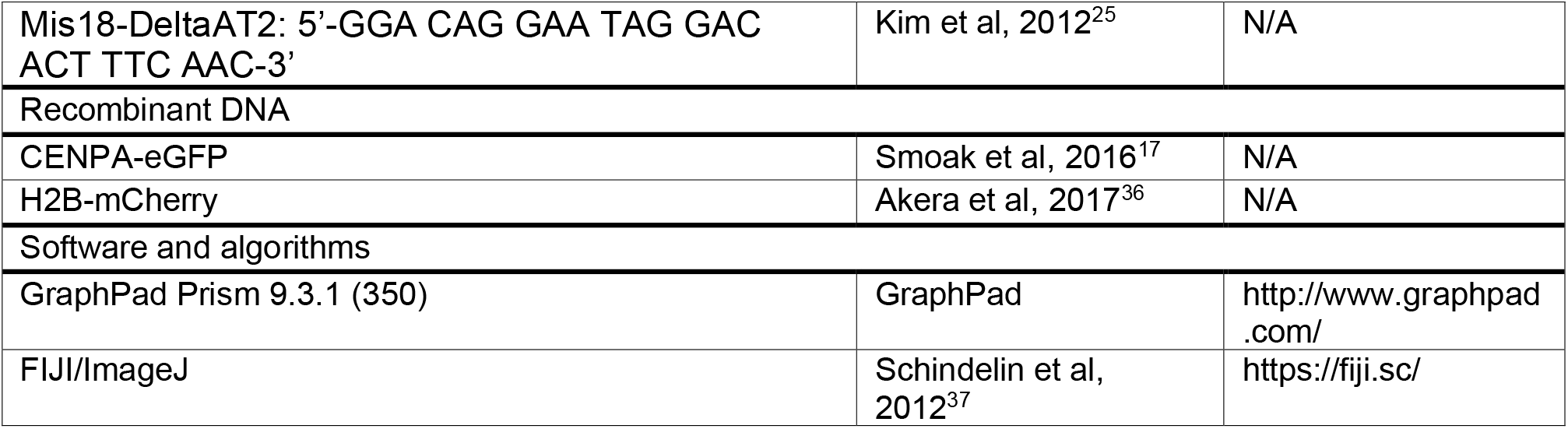

## RESOURCE AVAILABILITY

### Lead contact

Further information and requests for resources and reagents should be directed to and will be fulfilled by the lead contact, Ben E. Black.

### Materials availability

This study did not generate new unique reagents.

### Data and code availability

Images and measurement data reported in this paper will be shared by the lead contact upon request.

This paper does not report original code.

## METHOD DETAILS

### Animal husbandry and mouse strains

All animal experiments and protocols were approved by the Institutional Animal Use and Care Committee of the University of Pennsylvania and were consistent with National Institutes of Health guidelines (protocol #803994). Experimental animals were compared to age, gender and genetic background matched controls. The *Mis18a*^*fl/fl*^ strain is a previously published and validated knockout strain^25^. This strain is in a mixed genetic background of C57BL6/J/129Sv/CBA/J (generated by Sung Hee Baek, Seoul National University and obtained from Kikue Tachibana, Max Planck Institute). Oocyte specific conditional knockout strains *Mis18a*^*fl/fl*^*;Zp3-Cre* (*Mis18α* late KO) and *Mis18a*^*fl/fl*^*;Gdf9-iCre* (*Mis18α* early KO) were generated by crossing the original *Mis18a*^*fl/fl*^ strain to either *C57BL/6-Tg(Zp3-cre)93Knw/J* (RRID:IMSR_JAX:003651, Jackson Laboratory) or *Tg(Gdf9-icre)5092Coo/J* (RRID:IMSR_JAX:011062, Jackson Laboratory). The following primers were used to genotype the animals: 1) *Mis18a*^*fl/fl*^: SDL-Forward: TGC CTA TTG GTG TAC CTT CCA GTG, LOX-Reverse: CCT AAG TCG TTG ACC TGA CCG AGG, 2) *Mis18a^fl/Δ^* : Mis18-DeltaAT2 Reverse: GGA CAG GAA TAG GAC ACT TTC AAC combined with SDL-Forward (Figure S1).

### Oocyte collection, meiotic maturation, and culture

Female mice were hormonally primed with 5 U of pregnant mare serum gonadotropin (PMSG, Peptides International) 44-48 h before oocyte collection. Germinal vesicle intact oocytes were collected in bicarbonate-free minimal essential medium (M2, Sigma), denuded from cumulus cells, and cultured in Chatot–Ziomek–Bavister^38^ (CZB, Fisher Scientific) medium covered with mineral oil (Sigma, BioXTRA) in a humidified atmosphere of 5% CO_2_ in air at 37 °C. During collection, meiotic resumption was inhibited by addition of 2.5 mM milrinone (Sigma). Milrinone was subsequently washed out to allow meiotic resumption and oocytes were fixed 6–7 h later at metaphase I.

### Oocyte immunocytochemistry

Oocytes were fixed in 2% paraformaldehyde (Sigma) in phosphate buffered saline (PBS) with 0.1% Triton X-100 (Sigma), pH 7.4, for 20 min at room temperature (r.t.), permeabilized in PBS with 0.2% Triton X-100 for 15 min at r.t., placed in blocking solution (PBS containing 0.3% bovine serum albumin (BSA) and 0.01% Tween-20) for 20 minutes at r.t., treated with λ-phosphatase (1,600 U, NEB) for 1 h at 30 °C for CENP-A staining, incubated for 1 h with primary antibody in blocking solution, washed three times for 10 min each, incubated for 1 h with secondary antibody, washed three times for 10 min each, and mounted in Vectashield with 4′,6-diamidino-2-phenylindole (DAPI; Vector) to visualize the chromosomes. The primary antibody was rabbit anti-mouse CENP-A (1:200, Cell Signaling, C51A7). The secondary antibody was donkey anti-rabbit Alexa Fluor 488 (1:500, Invitrogen).

### Microscopy

Confocal images were collected as *z*-stacks with 0.5-μm intervals, using a microscope (DMI4000 B; Leica) equipped with a 63x 1.3-NA glycerol-immersion objective lens, an *x*–*y* piezo Z stage (Applied Scientific Instrumentation), a spinning disk confocal scanner (Yokogawa Corporation of America), an electron-multiplying charge-coupled device camera (ImageEM C9100-13; Hamamatsu Photonics) and either an LMM5 (Spectral Applied Research) or Versalase (Vortran Laser Technology) laser merge module, controlled by MetaMorph software (Molecular Devices, v7.10.3.294). Images were acquired using the same laser settings and all images in a panel were scaled the same. Single channels are shown wherever quantifications were performed.

### Quantification of centromere signals

To quantify centromere signal ratios, a sum intensity Z-projection was made using Fiji/ImageJ software. Circles of constant diameter were drawn around individual centromeres and the average intensity was calculated for each centromere after subtracting background, obtained from nearby regions. Raw centromere intensities were obtained from several controlled independent experiments and multiple cells were analyzed from each animal. Normalization of centromere intensities was performed using age- and gender-matched controls for each independent experiment.

### Embryo collection, microinjection and culture

*Mis18a*^*fl/fl*^ or *Mis18a*^*fl/fl*^*;Zp3-Cre* (*Mis18α* late KO) females were hormonally primed with 5 U of PMSG (Peptides International) and the oocytes were matured *in vivo* with 5 U of human chorionic gonadotropin (hCG; Sigma) before mating with B6D2F1/J males (F1 hybrid of a cross between C57BL6/J and DBA2/J; RRID:IMSR_JAX:100006, Jackson Laboratory). Males were fed a special low soymeal diet (5LG4 irradiated diet, Labdiet) and housed singly. Embryos were collected 14–16 h post hCG in M2 containing hyaluronidase (0.3 mg ml^−1^) to remove cumulus cells and subsequently washed in M2 (Sigma). and cultured in EmbryoMax Advanced KSOM (AKSOM, Millipore Sigma) with humidified air and 5% CO_2_. Embryos were then subjected to inter-cytoplasmic microinjection in M2 medium covered with mineral oil (Sigma, BioXtra) at r.t. with a micromanipulator (Narishige) and a picoinjector (Medical Systems Corp). Each embryo was injected with 2 pl of cRNA, then cultured in EmbryoMax Advanced KSOM (AKSOM, Millipore Sigma) with humidified air and 5% CO_2_ until 2 cell stage and fixed in 2% paraformaldehyde. The following cRNAs were used for microinjection: H2B-mCherry (human histone H2B with mCherry at the C-terminus) at 25 ng/ul, CENP-A-EGFP (mouse CENP-A with EGFP at the C-terminus) at 20 ng/ul. The cRNAs were synthesized using the T7 mScript Standard mRNA kit (Thermo Fisher Scientific) and purified by phenol-chloroform extraction.

## Data and code availability

All data are available upon request. No unique code was generated or used in this study.

## SUPPLEMENTAL FIGURES AND LEGENDS

**Figure S1:**
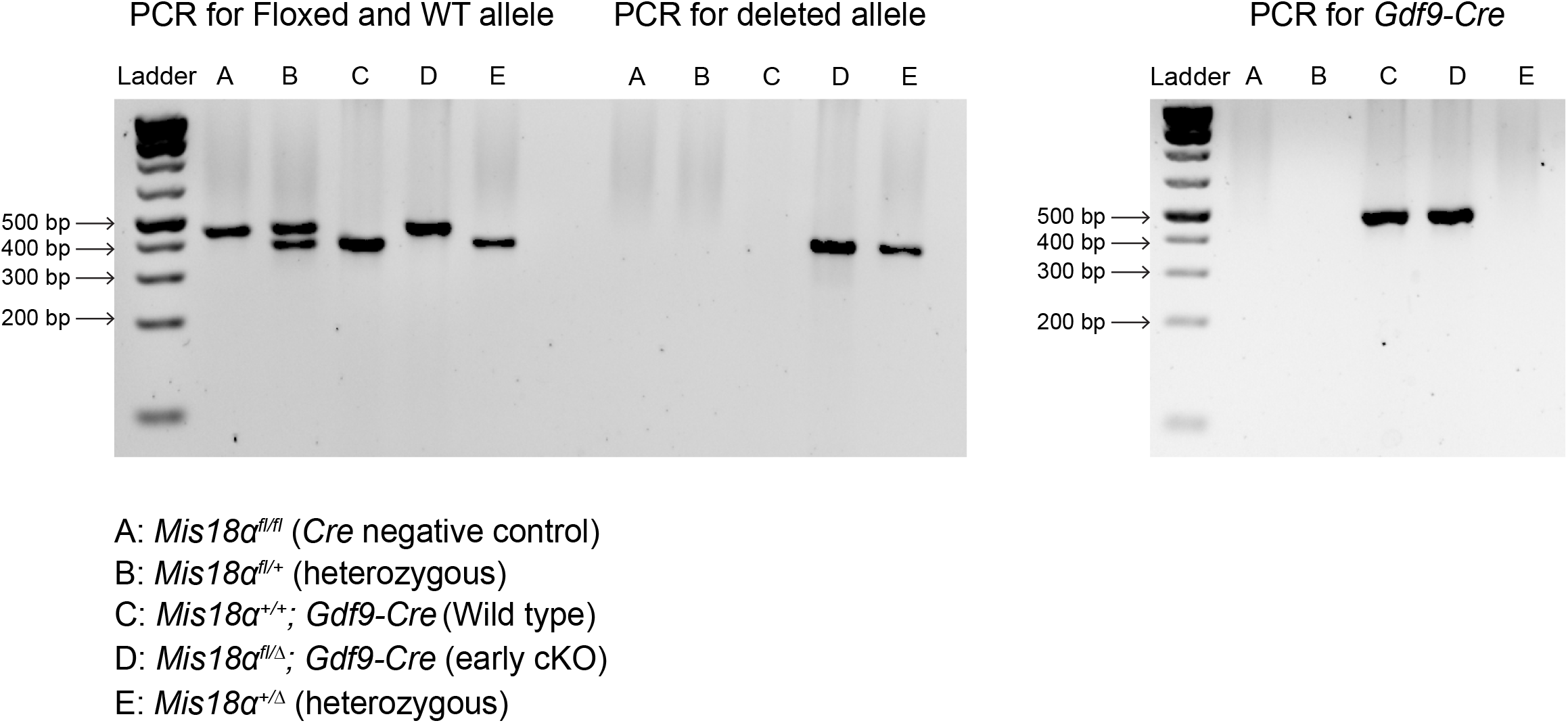
Genotyping gels for *Mis18α*^*fl/fl*^ mice. Related to Figure 1.

**Figure S2:**
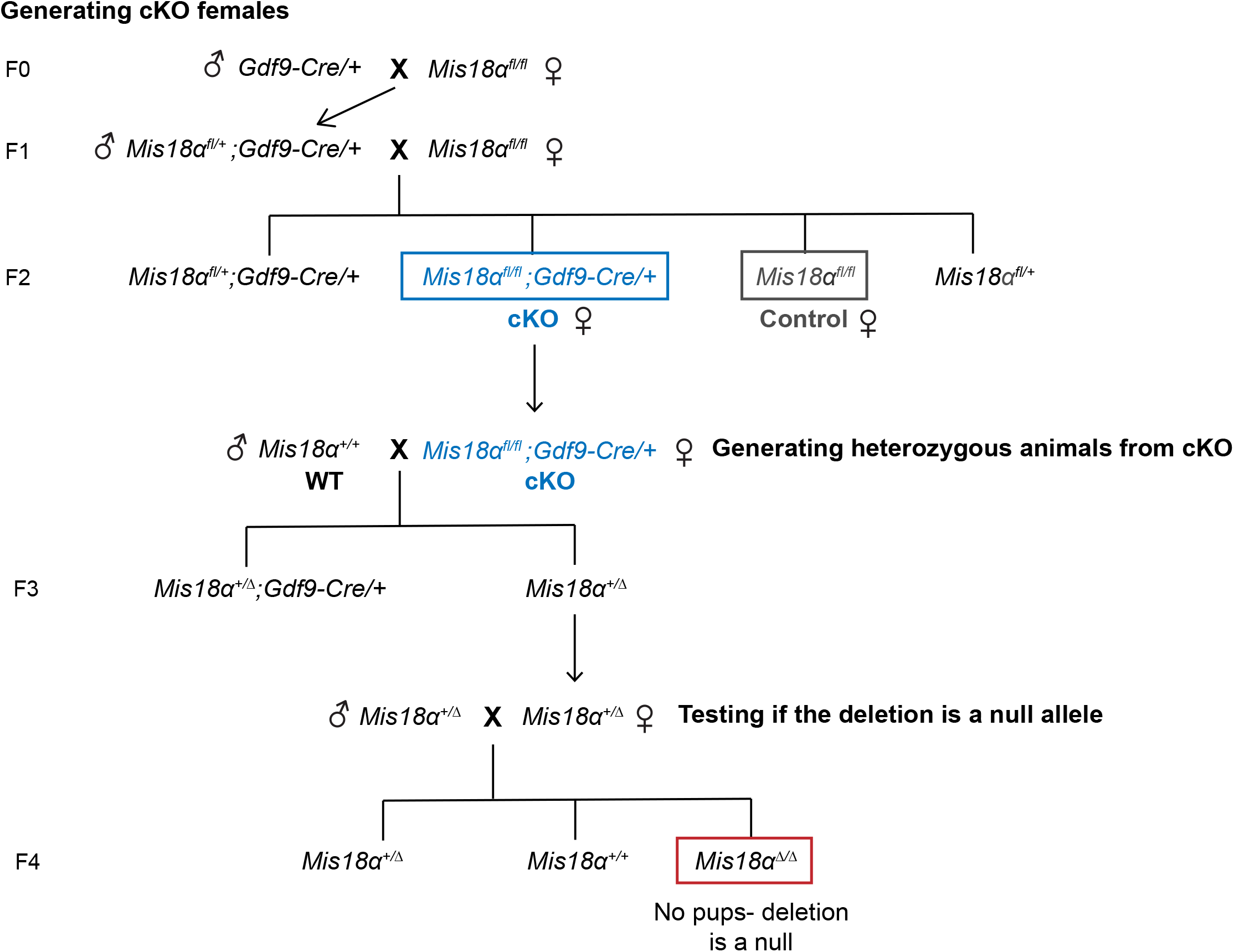
Cross scheme to combine the *Cre* recombinases with the *Mis18α* floxed allele. Related to Figures 1 and 2. Crosses are shown for *Gdf9-Cre*, and equivalent crosses were done for *Zp3-Cre*. The blue and grey boxes designate cKO and control females used for CENP-A measurements in oocytes (Figure 4). The cKO mothers were also used to generate heterozygous animals, which were then utilized to confirm that *Mis18α* deletion is a null allele (also see Figure 1D).

